# Molecular profiling predicts meningioma recurrence and reveals loss of DREAM complex repression in aggressive tumors

**DOI:** 10.1101/679480

**Authors:** Akash J. Patel, Ying-Wooi Wan, Rami Al-Ouran, Jean-Pierre Revelli, Maria F. Cardenas, Mazen Oneissi, Liu Xi, Ali Jalali, John F. Magnotti, Donna M. Muzny, HarshaVardhan Doddapaneni, Sherly Sebastian, Kent A. Heck, J. Clay Goodman, Shankar P. Gopinath, Zhandong Liu, Ganesh Rao, Sharon E. Plon, Daniel Yoshor, David A. Wheeler, Huda Y. Zoghbi, Tiemo J. Klisch

## Abstract

Meningiomas account for one-third of all primary brain tumors. Although typically benign, about 20% of meningiomas are aggressive, and despite the rigor of the current histopathological classification system, there remains considerable uncertainty in predicting tumor behavior. Here, we analyzed 160 tumors from all three WHO grades (I-III) using clinical, gene expression and sequencing data. Unsupervised clustering analysis identified three molecular types (A, B, and C) that reliably predicted recurrence. These groups did not directly correlate with the WHO grading system, which would classify more than half of the tumors in the most aggressive molecular type as benign. Transcriptional and biochemical analyses revealed that aggressive meningiomas involve loss of the repressor function of the DREAM complex, which results in cell cycle activation; only tumors in this category tend to recur after full resection. These findings should improve our ability to predict recurrence and develop targeted treatments for these clinically challenging tumors.

**Significance Statement:** Meningiomas are the most common primary brain tumors. Although most of these tumors are benign, one-fifth will recur despite apparently complete resection. Several studies have demonstrated that genomic approaches can yield important insights into the biology of these tumors. We performed RNA sequencing and whole-exome sequencing of 160 tumors from 140 patients, which identified three distinct groups of meningioma that correlate with recurrence better than the current WHO grading system. Our analysis also revealed that the most aggressive type was characterized by loss of the repressive DREAM complex. These findings should improve prognostication for patients and lead to viable therapeutic targets.

## Introduction

Meningiomas are the most common primary tumors of the brain and central nervous system (1, 2), and they are most commonly benign (WHO grade I). Nevertheless, roughly 20% of meningiomas are atypical (grade II) or malignant (grade III), with a five-year recurrence rate of up to 41% (3–5); such tumors require serial resections until they become inoperable, and the five-year survival rate can be as low as 35% (6). At present, the WHO histopathological classification system does not consistently predict whether an individual meningioma will recur after complete surgical resection (7). We clearly need a better understanding of meningioma biology in order to develop effective complements to surgery and radiation.

There are good reasons to believe that meningioma might be amenable to the sort of molecular profiling that has transformed the diagnosis and treatment of medulloblastoma, glioma, and many other cancers in recent years (8–11). The first hint of an underlying genetic mechanism came from the observation that meningiomas frequently arise in the context of neurofibromatosis type 2 (NF2) (12). In fact, half of sporadic meningiomas and a majority of higher-grade tumors involve loss of NF2 function or loss of heterozygosity of chromosome 22q, where *NF2* is located (13, 14). Several whole exome/genome sequencing studies have identified recurrent somatic mutations in *TRAF7, KLF4, AKT1, SMO*, and *POLR2A* in benign (grade I) tumors (15–17). Harmanci et al. found that a majority of primary atypical meningiomas have loss of *NF2* along with either genomic instability or *SMARCB1* mutations (13); this combination of features was not able to completely separate atypical from benign tumors, but the addition of the top 25 most differentially expressed genes raised the prediction accuracy of the model to 91% for atypical tumors with a high or medium Ki-67 index. Bi et al. found that grade III tumors are less likely to have *TRAF7*, *KL4*, *AKT1*, or *SMO* mutations but more likely to show genomic instability (copy number variation or CNV) (18). Vasudevan et al. sought targetable pathways in high-grade meningiomas and found that high *FOXM1* expression is associated with poor clinical outcomes (19); this is one of several studies showing that DNA methylation profiles have clinical relevance (14, 19–21).

All these studies demonstrate that molecular approaches yield important insights. Yet most relied on the existing WHO histopathological classification system, i.e., they studied tumors within specific WHO grades. To our knowledge, only Sahm et al. studied meningiomas across all grades, using methylation arrays to find two major epigenetic groups with six subclasses between them (14). Given that global epigenetic changes are just one mechanism by which cells alter expression of large groups of genes, we decided to focus on transcriptional profiling. This approach has the advantage of yielding functional biological information about tumor behavior. We therefore used an unsupervised approach to analyze RNA sequencing (RNA-seq) and whole-exome sequencing (WES) data from 160 fresh-frozen grade I, II and III meningioma samples. Our analysis yielded three distinct types of meningioma that correlate with clinical outcomes better than the WHO classification; it also revealed a molecular signature for the most aggressive tumors that provides biological insight into their etiology.

## Results

### Patient Demographics and Pathologic Characteristics

We analyzed 160 meningioma samples from 140 patients (see Methods for details). According to the WHO histopathological classification system for meningioma, 121 tumors were grade I (benign), 32 were grade II (atypical), and 7 were grade III (malignant). Female sex confers greater risk for meningioma (1), and our cohort reflected the expected proportions, with 90 (64%) female and 50 (36%) male subjects. The median age at the time of initial surgery for these patients was 60 years (range 21-81 years). Seventy-nine percent of patients underwent a gross total resection, twenty-two percent underwent a subtotal resection, and in one case the extent of resection was unknown. The follow-up period ranged from 0 to 91 months (median 28 months). Twenty-four tumors (17%) had a local recurrence. The recurrence rate for WHO I grade tumors was 11%; grade II, 42%; and grade III, 83%. The patient characteristics and pathology of our cohort are presented in Dataset S1. None of the tumors in our discovery or independent validation set had been treated with adjuvant radiation prior to profiling. Five patients had had radiation as children (four for cancers and one for tinea capitis); these are marked with an asterisk in Dataset S1.

### Identification of Meningioma Subtypes by Transcriptome Analysis

To determine whether meningiomas could be differentiated based on gene expression profiles, we used principal component analysis (PCA) on a discovery set of 97 tumors (77 WHO grade I, 20 WHO grade II; of note, we had no primary grade III tumors, which are exceedingly rare as they are usually recurrences (22)). The tumors did not cluster into distinct groups based on WHO grade (Fig. 1A).

**Fig. 1.**
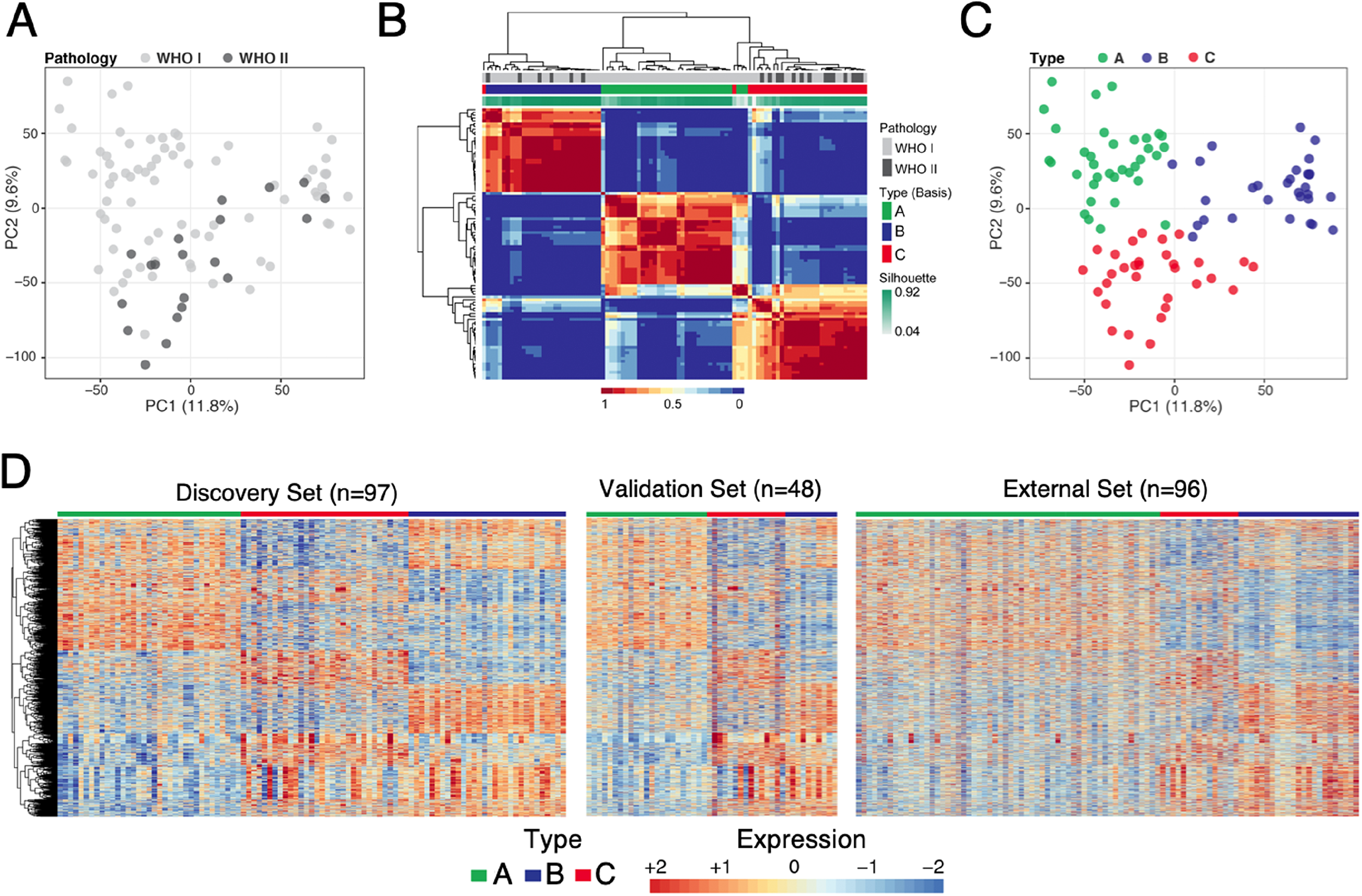
Identification of meningioma subtypes using gene expression profiles. (**A**) Principal component analysis (PCA) on all genes of 97 tumors colored by WHO grading. WHO grade I tumors are represented by light grey circles; WHO grade II tumors are represented by dark grey circles. (**B**) Consensus matrix of the tumors for k=3 from 1000 runs of non-negative matrix factorization analysis depicts three distinguishable types based on the gene expression data. (**C**) PCA on all genes, colored according to molecularly-defined types. (**D**) Expression heatmap of the 3,484 genes common in all three data sets. The expression patterns of these genes distinguish three expression types in our discovery set (left panel), validation set (middle panel) and publicly available data set (right panel). Type A is labelled in green; type B, blue; type C, red. Labels in the two independent validated set were predicted using a random forest trained on discovery set.

We then employed non-negative matrix factorization (NMF) clustering for k=2 to k=7 using the 1,500 genes that varied most among the tumor samples. NMF is an unsupervised machine learning approach commonly used for cancer subtypes discovery (8). After 1,000 iterations, three clusters (k=3) emerged as providing the best fit as determined by the consensus membership, cophenetic and silhouette scores (Fig. 1B, SI Appendix Fig. S1A-C). We evaluated the cluster significance of the three subtypes using SigClust (23), and observed statistical significance between cluster boundaries (SI Appendix Fig. S1D). The three clusters can also be discerned from the expression heatmap (SI Appendix Fig. S1E) and exhibit significant differences in WHO grade representation (p-value = 0.0020, ANOVA): A (green) is populated exclusively with WHO grade I tumors; B (blue) contains mostly WHO grade I (79%) tumors, with 21% grade II; and C (red) contains similar proportions of WHO grade I and II tumors (56% and 44%, respectively; Dataset S1). Because the WHO grade III tumors in our cohort were all recurrences, they were not included in the primary transcriptome analysis.

To understand the robustness of the three molecular subtypes, we examined the gene expression profiles associated with each cluster in two independent data sets: an independent cohort of 48 tumors (39 WHO grade I and 9 WHO grade II) and a published microarray dataset of 96 meningiomas (16). Since the three data sets were profiled on two different platforms, we first filtered out genes that are not expressed in any tumors across the three datasets. Then, on the discovery data set, we performed pairwise comparisons between each cluster to identify genes that are differentially expressed with a minimum absolute fold-change of 1.5 and a false discovery rate of 1%. This yielded 3,484 genes, which we used to build a Random Forest classifier to predict a cluster label of each sample. The Random Forest was trained on the discovery data set of 97 samples. To avoid overfitting due to small sample size, we evaluated the fitted model in two independent validation data sets which were never used for feature selection or training of our random forest calssifier. We observed concordant gene expression patterns for all three clusters across training and validation sets (Fig. 1D). These results provide evidence that the molecular types designated by differential gene expression of our discovery set are stable, even across platforms.

We next analyzed the association of clinical variables with the three transcriptionally defined types (from here we will refer to our three molecular classes as types A, B, and C, as distinct from the WHO classification system’s grades I, II, and III). One important clinical variable is the MIB1 index, a measure of the mitotic activity of the tumor, which has prognostic significance (24). The median MIB1 index tracked with molecular type, being lowest in type A, intermediate in type B, and highest in type C (2.5, 3.5, and 6.3, respectively; Fig. 2A upper panel, p-value = 0.0026, ANOVA), despite 56% of our type C tumors being classified as WHO grade I. While the sample size is smaller in our validation cohort, we observed the same trend (Fig. 2A, lower panel). To ensure that these differences were not due to a mixture of WHO grade tumors in the types, we analyzed the MIB1 index of only the WHO grade I tumors (Fig. 2B, left panels) and only the grade II tumors (Fig. 2B, right panels) and found the same tracking of MIB1 index from molecular types A to C, within each WHO grade. Because the MIB1 index is based on Ki67 immunohistochemistry staining, which is subject to interobserver variability (25, 26), we quantified its average transcript levels (MKI67) and observed a concordant result: statistically significant increases from molecular types A to C in our discovery cohort (SI Appendix Fig. S2A, upper panel; p-value < 0.0001) and an identical trend in our validation cohort (SI Appendix Fig. S2A, lower panel).

**Fig. 2.**
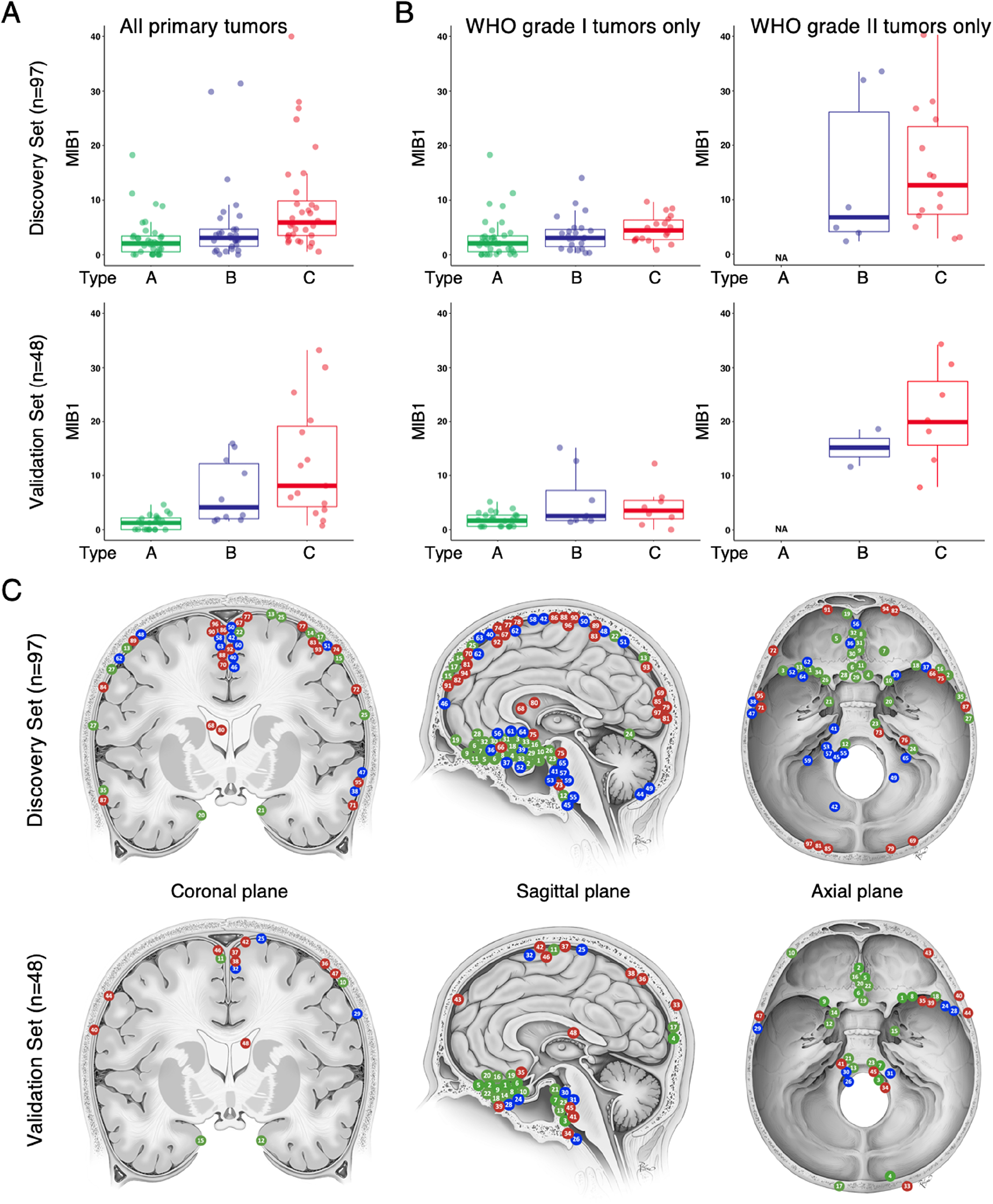
Clinical characteristics of gene expression-defined meningioma types. Type A is labelled in green; type B, blue; type C, red. (**A**) Boxplots showing the median MIB-1 index for types A to C in the discovery set (p-value=0.0026, ANOVA, upper panel) and the validation set (p-value<0.0001, ANOVA, lower panel). (**B**) Boxplot of the MIB1 for types A to C for only WHO grade I tumors (left panels) in the discovery set (p-value=0.4359, ANOVA, upper panel) and the validation set (p-value=0.0044, ANOVA, lower panel) and WHO grade II tumors (right panels) in the discovery set (p-value=0.6059, ANOVA, upper panel) and the validation set (p-value=0.3380, ANOVA, lower panel). (**C**) Location of tumors in our cohort in the discovery set (upper panels) and validation set (lower panels). Each tumor is marked on 2 views, either coronal and sagittal or axial and sagittal, respectively (see Dataset S1).

It has been reported that tumors with different somatic mutations cluster to different intracranial regions (e.g., TRAF7 and SMO mutant tumors tend to form in the anterior skull base) (15). We therefore asked whether any of our three molecularly-defined types were associated with specific locations (Fig. 2C and Dataset S1). Although the sample size relative to the number of factors precludes making strong conclusions, we used a generalized linear model (Poisson link function) analysis to compare the distribution of tumors for each type across 16 anatomical locations. Only the anterior skull base and occipital locations showed a significant difference between types (adjusted p-value = 0.0004 and 0.0329, respectively), with type A tumors more likely to be located in the anterior skull base and type C tumors more likely to arise in the occipital region (Fig. 2C). We next compared the spatial distribution of tumor types between datasets (BIC comparison); we found no significant interactions with dataset (discovery vs validation) suggesting these patterns were consistent in both samples.

We also examined the gender distribution in our expression types. In our discovery set, types A and B show the expected 2:1 female to male distribution, but 56% of patients in type C are male (p-value = 0.0240, Dataset S1).

Lastly, we assessed the recurrence-free survival (RFS) across the three types (Fig. 3). We did not analyze overall survival because only three patients died (one in the discovery set and two in our validation set). We first analyzed our discovery set of 97 tumors. WHO grade II tumors tended to have a shorter RFS than WHO grade I tumors, but this trend did not quite reach significance (Fig. 3A left panel; log-rank p-value = 0.0530). On the other hand, our type A and B tumors had an indistinguishably long RFS with only four recurrences, even though 21% of the type B tumors would be classified by the current WHO system as “high grade.” Type C tumors, however, have a significantly shorter RFS than the two other types (Fig. 3A right panel; log-rank p-value < 0.0001), despite the fact that the majority of the tumors in type C are classified as WHO grade I.

**Fig. 3.**
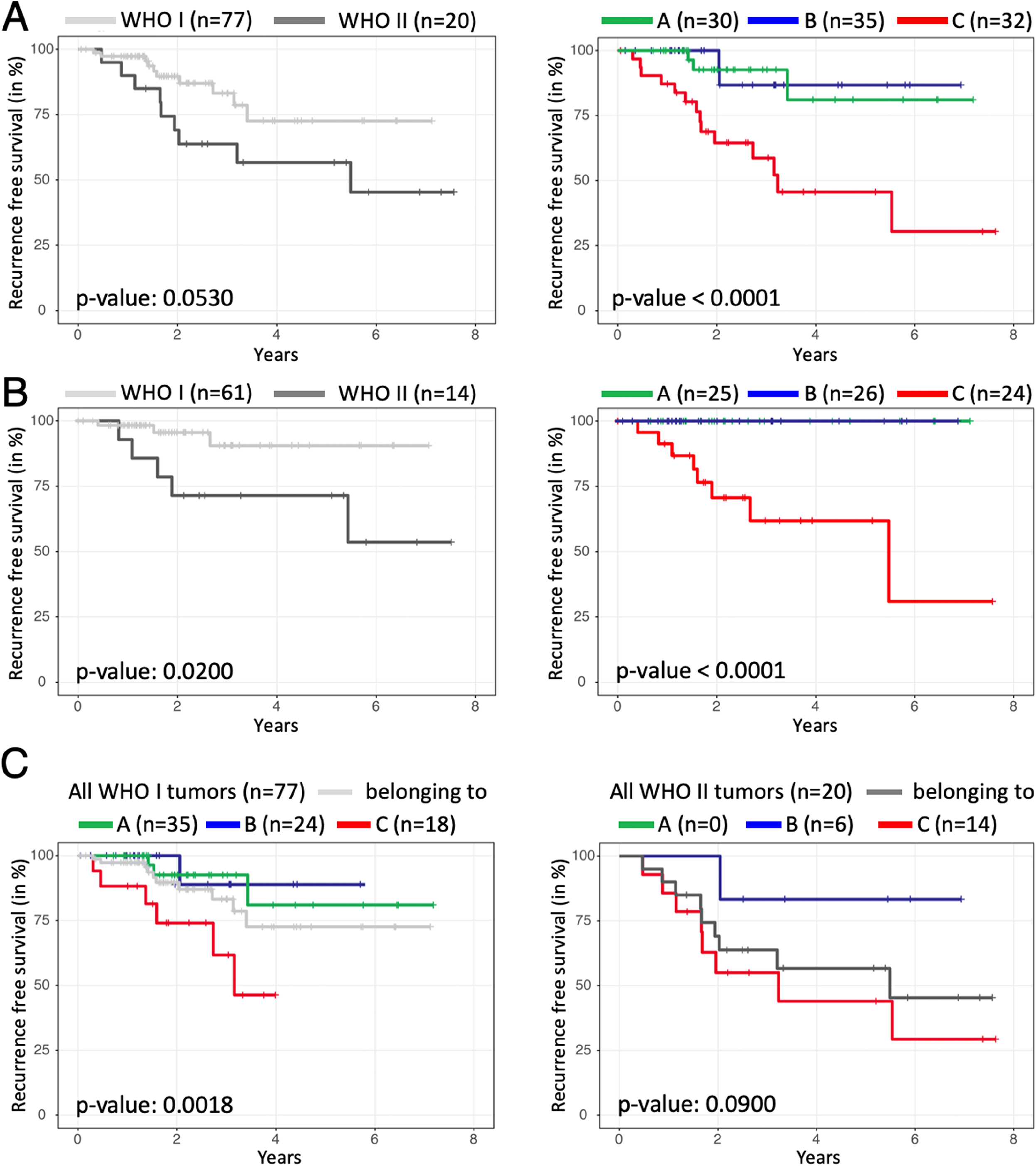
Recurrence-free Survival of WHO grade and gene expression-defined meningioma types. Recurrence-free survival analysis (RFS) based on (**A**) WHO grading (left panel) and by expression-defined types (right panel) in all tumors and (**B**) only tumors that underwent complete resection. (**C**) RFS for expression-defined types within only WHO grade I tumors (left panel) or WHO grade II tumors (right panel) shows the ability of the molecular typology to refine RFS despite WHO grading. N represents the initial number of tumors for each curve.

To ensure that the extent of resection was not responsible for these RFS differences, we looked at RFS for only those tumors that underwent gross-total resection. Both WHO grade I and II had recurrences, with more for WHO grade II (Fig. 3B, left panel, log-rank p-value = 0.0200), but of our molecularly defined types, only type C tumors recurred (Fig. 3B, right panel, log-rank p-value < 0.0001).

To rule out the effect of WHO grade on the recurrence trends seen with our types, we analyzed the RFS of our types within each WHO grade in our discovery cohort (similar to our MIB1 analysis). We found that type C WHO grade I tumors have much worse recurrence rates (33%) than type A (8.6%) or type B (4.2%) WHO grade I tumors or WHO grade I tumors as a whole (13%) (Fig. 3C, right panel; log-rank p-value = 0.0018, ANOVA). The same holds true within WHO grade II tumors (Fig. 3C, left panel; log-rank p-value = 0.0900, ANOVA): type C WHO grade II tumors have a 57% recurrence rate, higher than type B (16.7% recurrence rate) or all WHO grade II tumors (45% recurrence rate).

Thus, our expression-based classification identifies WHO grade I/II tumors that have a high risk of recurrence. These data also suggest that total resection is less likely to cure type C tumors.

### Copy Number and Somatic Alterations in Meningiomas

Since high-grade meningiomas have more chromosomal abnormalities (13, 18), we analyzed the three types of tumors for genomic instability using copy number data derived from whole-exome sequencing (WES) (Fig. 4). We had copy number data for 84 tumors in the discovery cohort and 44 in the validation cohort. Type A had no notable chromosomal losses or gains. Type B tumors showed significant loss of chromosome 22q, the most commonly reported chromosomal abnormality in meningioma (27) (Fig. 4A; 84%; p-value < 0.0001, Chi-square). Type C manifested the most genomic instability, showing loss of chromosome 22q (89%; p-value < 0.0001 Chi-square) and chromosome 1p (79%; p-value < 0.0001, Chi-square), the second most common reported abnormality (27, 28). Furthermore, over 20% of the type C tumors showed losses in chr3p, chr4p, chr6q, chr14p, chr14q, or chr18q (Fig. 4A upper panel). Both validation sets replicated these results (Fig. 4A, lower panels). Interestingly, the type of chromosome loss is almost sufficient to distinguish between types. Combining all data sets, retaining both chr1p and chr22q identified type A tumors with a 94% sensitivity and 86% specificity with a positive predictive value (PPV) of 86% and a negative predictive value (NPV) of 94%. Deletion of only chr22q identified type B with a sensitivity of 76%, specificity of 95%, PPV of 83% and NPV of 92%. Deletion of both chr22q and chr1p identified type C with a sensitivity of 68%, specificity of 95%, PPV of 83% and NPV of 90%. We also examined the distribution of chromosome loss by gender and did not find a significant difference between females and males (Fig. 4B). However, we noticed that the ratio of female to male differed between types: combining all data sets, type A has 78% female patients, type B has 77%, and type C only 51% (p-value = 0.0008, Chi-square).

**Fig. 4.**
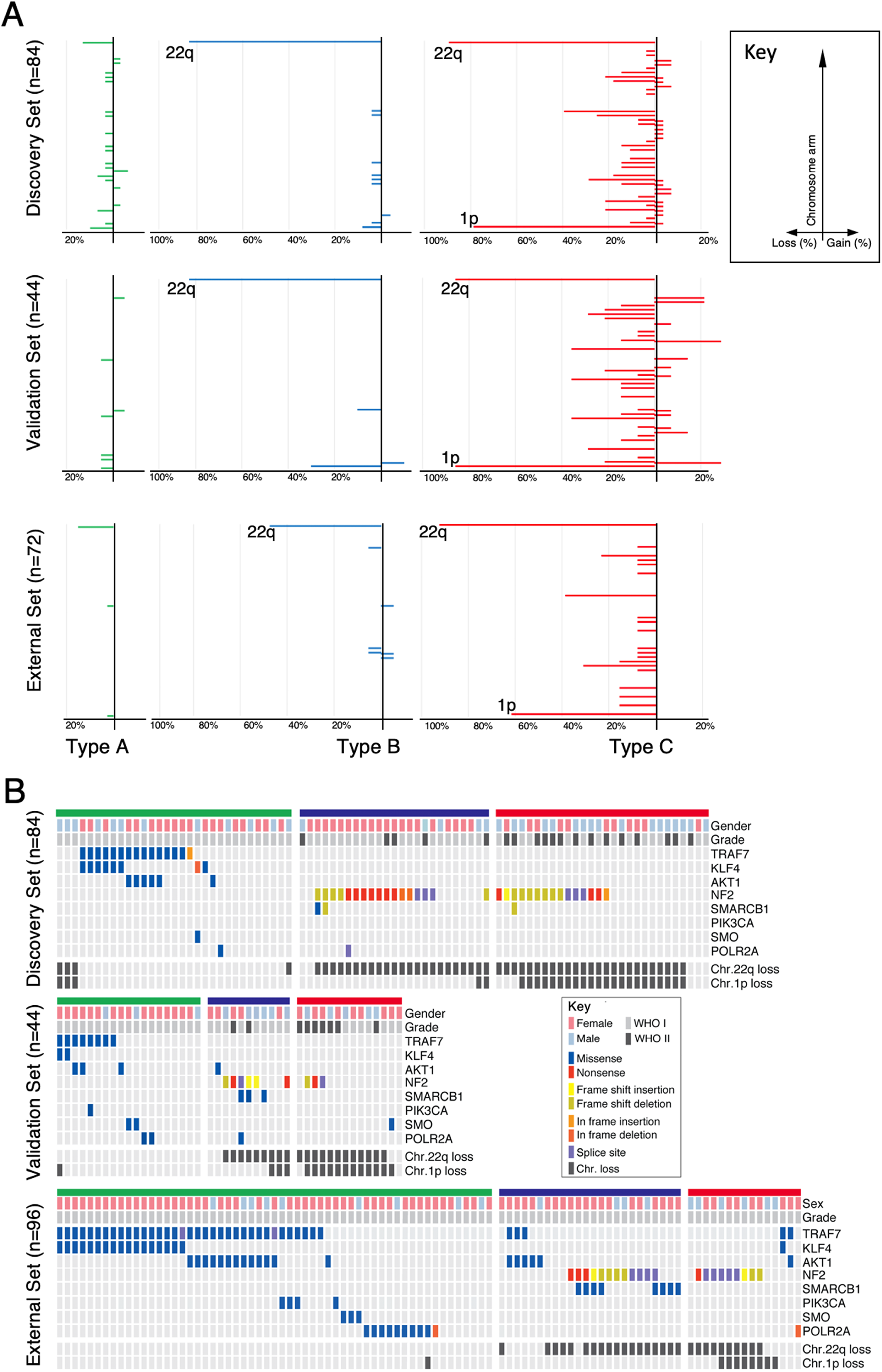
Genomic landscape of meningiomas by gene expression-defined types. Type A is labelled in green; type B, blue; type C, red. (**A**) Differences in chromosomal alterations by type are shown with losses to the left and gains to the right. (**B**) Oncoprint depicting the mutation profiles of each meningioma type in the discovery set (upper panel) the internal (middle panel) and external validation set (lower panel).

Given that the tumor types could nearly be distinguished based on copy number data alone, we calculated the PPV and NPV for recurrence based on copy number alterations. Loss of both chr22q and chr1p predicted recurrence with a sensitivity of 75%, specificity of 78%, PPV of 44% and NPV of 93%. For tumors that underwent complete resection in our cohort, loss of both chr22q and chr1p predicted recurrence with a sensitivity of 100%, specificity of 76%, PPV of 36% and NPV of 100%.

WES also revealed 3,094 somatic mutations in our discovery cohort with a median of 0.47 mutations per megabase, which did not differ between tumor types (type A, 0.44; type B, 0.40; type C, 0.52; p-value = 0.4951). Specific mutations did, however, cluster according to type. Only type A tumors contained mutations in *TRAF7* (Fig. 4B; 43%, 40% and 61% in the discovery, validation and external set, respectively). Type A also contained the highest percentage of *KLF4* (26%, 10%, 30%) and *AKT1* (19%, 15%, 23%) mutations (Fig. 4B). In contrast, NF2 mutations were seen only in types B (68%, 50%, 50%) and C (54%, 21%, 60%). Strikingly, these mutations were usually combined with a loss of the other allele on chromosome 22p. SMARCB1 mutations were primarily seen in type B, especially in the external set. *TERT* promoter mutations have been found in 13% of meningiomas and portend a worse prognosis (29–31). In both our discovery and validation set, there were 108 tumors whose sequencing included the *TERT* promoter. Of these, 13 tumors had a mutation in the *TERT* promoter, but they fell into all three tumor types (Dataset S1, p-value = 0.7623, Chi-square).

As the prevalence of NF2 mutations did not differ between types B and C (68% and 54%, respectively; p-value = 0.2837, Chi-square), we next explored whether the degree of NF2 expression loss could distinguish tumors in these types. Both types have markedly reduced levels of NF2 expression compared to type A (SI Appendix Fig. S2B, p-value < 0.0001) but did not differ from one another in this regard (SI Appendix Fig. S2B, p-value = 0.1400). Both showed typical loss of function variants (nonsense and frameshift) spanning the NF2 coding region (Fig. 4B and Dataset S2).

In sum, type A is characterized by recurrent somatic mutations in *TRAF7, KLF4* and *AKT1* but lacks any significant chromosomal gains/losses. Type B is characterized primarily by mutation in NF2 and loss of chr22q, and type C meningiomas have a significant burden of chromosomal gains/losses, most commonly loss of chr22q and chr1p together. Like WHO grade II and III tumors (22), our type C has a roughly equal proportion of females and males.

### Gene Set Enrichment Analysis Further Distinguishes Types B and C

To better differentiate types B and C and understand the biological pathways underlying these transcriptional changes, we performed gene set enrichment analysis (GSEA) (32, 33) for each expression type using the genes highly expressed in that type (Dataset S3). No single underlying pathway emerged for type A (Dataset S4). Four out of the five enriched categories in type B suggest that these tumors have lost the repressive activity of the PRC2 methyltransferase complex (Dataset S4). Genes highly overexpressed only in type C clustered in cell cycle modules, especially the G2/M checkpoint, which is regulated by the repressive transcription factors of the E2F family, such as E2F4, and its associated repressor, the DREAM complex (34). The two modules, “genes with promoters bound by E2F4” and “targets of the DREAM complex” were the most enriched modules (Dataset S4).

To determine whether these two repressor complexes truly reflect biological differences between these tumor types, we evaluated the enrichment scores of their target genes in all three types (Fig.s 5A). Strikingly, type B is characterized by the loss or dysfunction of the repressive PRC2 complex, whereas type C is characterized by loss or dysfunction of the repressive DREAM complex.

**Fig. 5.**
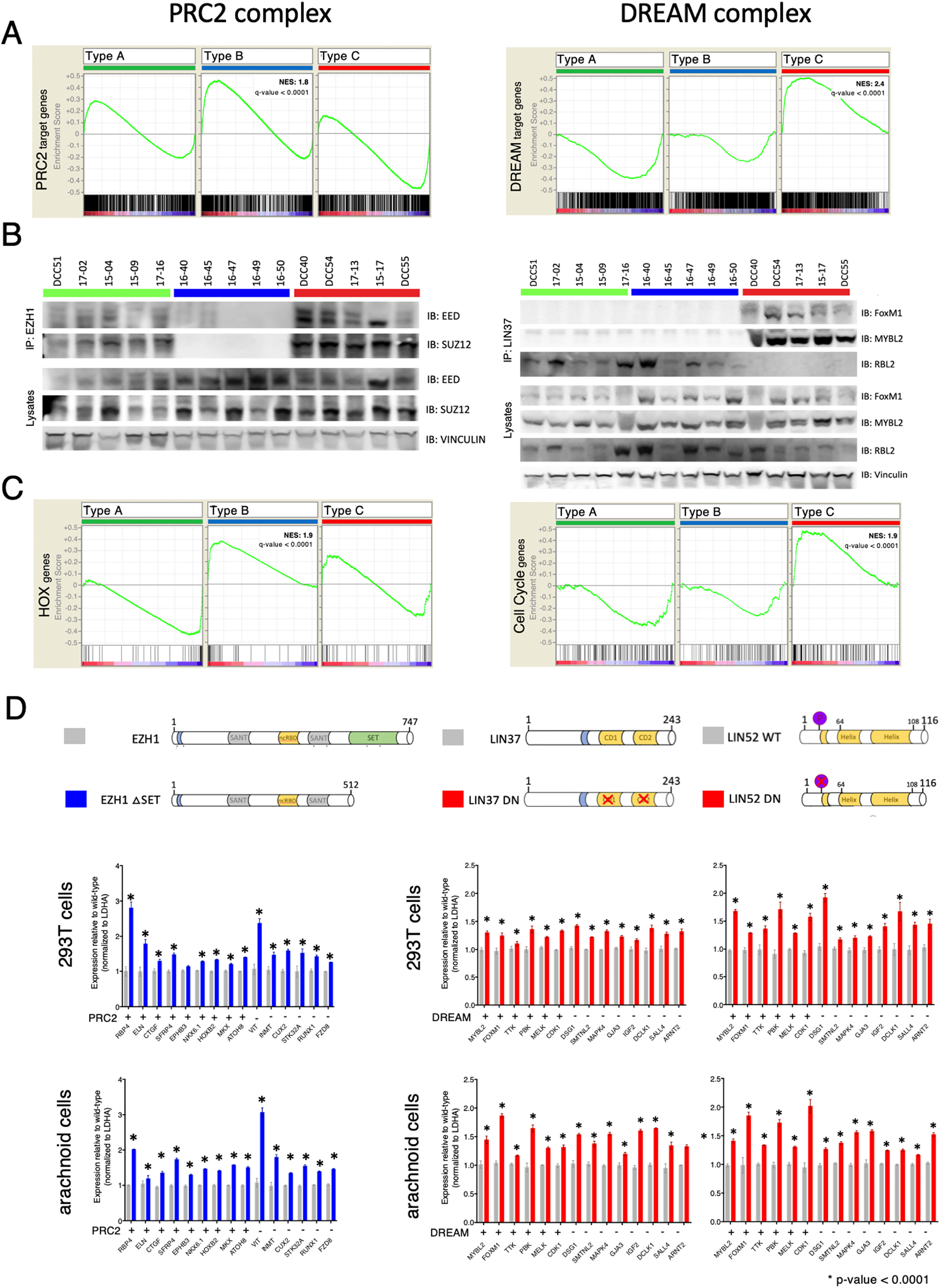
Validation of PRC2 and DREAM complex disruption in type B and C tumors, respectively. (**A**) GSEA analysis of the PRC2 (left) and DREAM (right) target genes from each type. (**B**) *Left*: Co-immunoprecipitation studies using 5 tumors per type for EZH1 then probed for anti-EED and anti-SUZ12. *Right*: Co-immunoprecipitation studies using 5 tumors per type for LIN37 then probed for anti-FoxM1, anti-MYBL2, and anti-RBL2. (**C**) GSEA analysis shows that HOX genes are enriched in type B (left) and cell cycle genes in type C (right). (**D**) qRT-PCR analysis measuring expression levels of type-specific upregulated genes and in 293T and arachnoid cells. *Left*: cells were transfected with either wild-type hEZH1 or hEZH1 ΔSET (dominant-negative EZH1). *Right*: cells were transfected with either wild-type hLIN37 or dominant-negative hLIN37 (left side) or either wild-type hLIN52 or dominant-negative hLIN52 (right side).

### Loss of the PRC2 Complex in Type B

The PRC2 complex is responsible for H3K27 di- and trimethylation and subsequent chromatin silencing. The core subunit consists of EED, SUZ12 and EZH1 or EZH2. We hypothesized that this complex is not forming or functioning in type B tumors, resulting in upregulation of the PRC2 target genes, as identified by the unbiased expression clustering. Therefore, we used cellular lysates from five tumors of each type, and immunoprecipitated the PRC2 complex using EZH1 (Fig. 5B). All tested proteins were expressed in all tumors (Fig. 5B, lysate lanes). Both EED and SUZ12 were detected in the EZH1 immunoprecipitates of type A and C tumors, but not type B tumors. This strongly suggests that the core complex is formed in type A and C tumors but not in type B tumors. Consistent with this finding, the PRC2 complex’s direct targets, the HOX transcription factors (35, 36), were significantly enriched only in type B (Fig. 5C; q-value < 0.0001).

To clarify whether loss of the PRC2 complex underlies the transcriptional dysregulation seen in type B, we transfected 293T cells with either wildtype EZH1 (1-747aa) or SET domain– depleted EZH1 (37) (EZH1-∆SET, 1-512aa, Fig. 5D). The SET domain of EZH1 is responsible for the lysine-specific histone methyltransferase activity: without this domain, PRC2 cannot perform H3K27 methylation, and so it enables aberrant gene activation (38, 39). After 48 hours of over-expression, we performed qRT-PCR analysis of fifteen genes that were all significantly upregulated in type B. We chose to include known PRC2 target genes (RBP4, ELN, CTGF, SFRP4, EPHB3, ATOH8) (40), including those which are also homeobox genes (NKX6.1, HOXB2, MKX) (40) (Dataset S3). We found all except one of these genes significantly upregulated in cells overexpressing EZH1-∆SET compared to wild-type EZH1 (Fig. 5D). Because meningiomas are thought to arise from arachnoid cap cells, we also generated an immortalized cell line from arachnoid cells (see methods for cell line establishment). As with the 293T cell line, the tested genes were upregulated upon loss of PRC2 complex function (Fig. 5D).

In sum, type B meningioma appear to have lost PRC2 complex function.

### Loss of the DREAM Complex in Type C

The DREAM complex is a highly-conserved master regulator of the cell cycle (34). It consists of MuvB core proteins: LIN52, LIN9, LIN37, LIN54 and RBBP4. When this core is bound to RB-like proteins (RBL1/2) and E2F, it forms the repressive DREAM complex, which keeps the cell quiescent. When the core associates with MYBL2 and FOXM1, however, it forms the activating DREAM complex, which allows cell cycle progression and subsequent proliferation. Interestingly, tumors from type C had the highest proliferation index, and a recent study found elevated expression of FOXM1 associated with high-grade meningiomas (13, 19).

We found increased expression of both FOXM1 and MYBL2 in our type C tumors, which aligns with our previous results suggesting that the DREAM complex has lost its repressive activity and allowed upregulation of these two target genes. To confirm that type B and type C tumors differ in the form of the DREAM complex that they express, we immunoprecipitated the core complex in tumors from all three types using LIN37. All investigated proteins were expressed in all tumors (Fig. 5B, lysate lanes), but RBL2 was associated with the core only in type A and B tumors (Fig. 5B). On the other hand, only in type C tumors was the core associated with both FOXM1 and MYBL2. Thus types A and B contain the repressive form of the DREAM complex, whereas type C tumors contain the activator forms of the complex.

If this is indeed the case, we would expect to see increased expression of DREAM target genes in type C tumors and decreased levels in the other two types. GSEA revealed that known DREAM target genes are highly enriched only in type C tumors (Fig. 5C; q-value < 0.0001). Using a strategy similar to that used for type B and the PRC2 complex, we took advantage of recently published dominant-negative forms of two MuvB core members, LIN37 (41) and LIN52 (42). It has been shown that mutations in two small domains in LIN37 (CD1 and CD2) result in the loss of the repressive function of the DREAM complex (LIN37-WT (1-243aa) and LIN37- DN) (41), and inhibiting phosphorylation of Serine28 on LIN52 results in similar phenotypes (LIN52-WT (1-116) and LIN52-DN) (42). We performed qRT-PCR analysis of fourteen genes that were all significantly upregulated in type C, some of which were known DREAM targets (MYBL2, FOXM1, TTK, PBK, MELK, and CDK1) (43). Overexpression of both dominant-negative constructs resulted in the upregulation of these genes in both 293T cells (Fig. 5D) and our arachnoid cell line (Fig. 5D).

In sum, type C meningiomas appear to be characterized by loss of the repressive DREAM complex function.

### Recurrent Tumors Match the Gene Expression Profile of the Original Tumor

Nine patients in our cohort had at least one resected recurrence with tissue available; two of these patients had multiple recurrences. Tumor progression was seen in three patients with type A tumors who could not have a complete resection due to tumor location. Gene expression profiling of these recurrent tumors demonstrated that they all remained within the type of the original tumor (Fig. 6, Dataset S1). Interestingly, three patients started with WHO grade II tumors that progressed to WHO grade III tumors, but their transcriptomic classification nonetheless remained the same (type C). Similarly, three patients with WHO grade I tumors had inexplicably rapid recurrences, despite complete resection in one; both their primary and recurrent tumors were type C. Of note, one recent study showed that a single grade III meningioma can be genetically heterogeneous (31); while we did not transcriptionally profile different parts of a tumor, this is something to be considered in future studies. Nevertheless, the consistency of the recurrent tumor profiles with that of their primary tumors suggests that the type C classification remains stable over time.

**Fig. 6.**
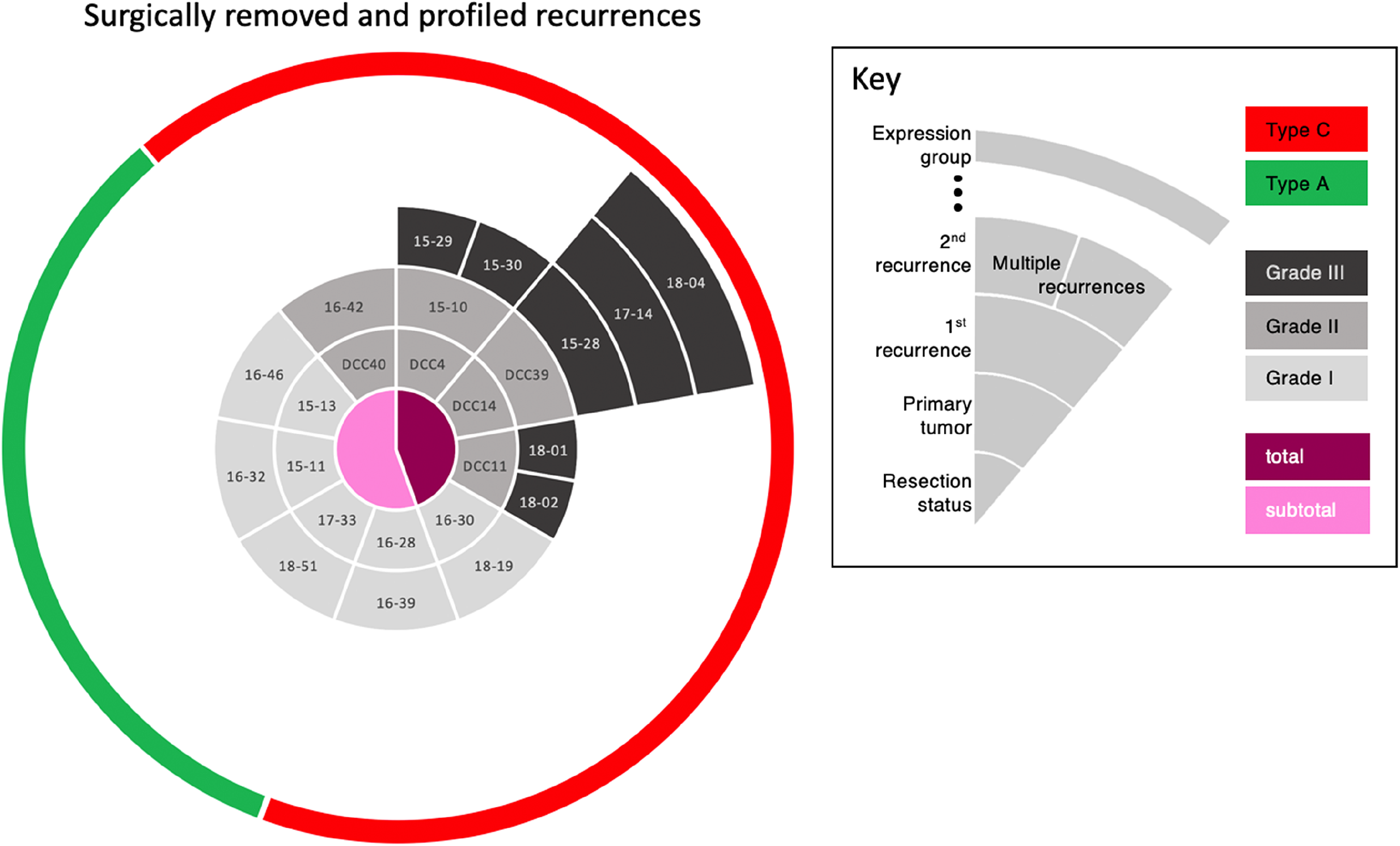
Timeline of tumor recurrences by patient. The central circle indicates whether the resection of the primary tumor was partial (pink) or total (magenta). The second circle identifies the primary tumor by ID number (light gray represents WHO grade I, medium grey is grade II). The third circle identifies the first recurrence, with tone of grey indicating the WHO grade. The remaining segments, all dark grey (grade III), indicate further recurrences over time. The green and red in the rim of the circle denote types A and C, respectively. Strikingly, whereas the WHO classification changes from primary tumor to recurrence in many cases (different shades of grey), the transcriptomic type classification did not—a primary tumor of type C remained type C throughout all recurrences.

## Discussion

The current histopathologic system for classifying meningiomas has shown some ability to predict clinical course, with WHO grade II and III tumors generally tending to recur. Yet a substantial number of grade I tumors also recur, despite successful resection and apparently benign features. Here we used unsupervised gene expression clustering of RNA-seq data from a large cohort of meningioma tumors to define new types that better correlate with recurrence-free survival and proliferation as measured by the MIB1 index. Most importantly, our expression-based model identifies tumors that are at high risk for recurrence, including those that would be classified as WHO grade I. For example, one of the patients in our clinic whose tumors are profiled in this study had total resection of a grade I tumor that recurred two years later; complete resection of this second tumor, also grade I, was insufficient to prevent a second recurrence 18 months later—still grade I, although the MIB1 index had risen to 9.1. Our molecular classification, however, identified all the tumors in this patient as type C.

A major difference between our study and previous explorations of the genomic landscape of meningioma is that those studies maintained the framework of the existing WHO histopathological classification for their molecular analysis (13, 15–19, 44). For example, one study of atypical (grade II) tumors found the majority to have NF2/chr22q loss and genomic instability along with overexpression of the E2F2 and FOXM1 transcriptional networks (13). Another study of high-grade tumors found overexpression of FOXM1 to be associated with poor clinical outcome (19). Only by analyzing both low and high-grade tumors in our cohort were we able to distinguish the different biology of type B and C tumors.

There are two limitations to our study. Several previous studies have explored DNA methylation profiling (14, 19–21), and it would be ideal to compare our classification system with that proposed by Sahm et al. by performing RNA-seq and methylation profiling on the same samples (14). (Methylation and expression may (45) or may not (46) correlate.) Unfortunately, the data in these papers are not publicly available, and thus could not be compared. The second limitation is the relatively short median follow-up of 28 months for tumors with a typically indolent course. Even so, our recurrence-free survival data were sufficient to identify an aggressive type of meningioma that would otherwise be classified as WHO grade I.

### Salient features of types A, B, and C

In our cohort, type A tumors were characterized by mutations in *TRAF7*, *KLF4*, and *AKT,* which confirms previous observations in benign meningiomas (15, 17). It is possible that the downstream consequences of these mutations converge biologically.

In our type B tumors, 91% showed loss of chromosome 22q. Our data suggest that these tumors lose PRC2 complex function and recur as infrequently as type A tumors. (A handful of type A tumors also had chr22q loss, but the number was only within the expected error rate based on the size of our discovery cohort.) Our current data suggest that if type A and B tumors are completely resected, they do not recur.

Type C tumors were instead characterized by the activator forms of the DREAM complex and subsequent upregulation of its target genes, including MYBL2 and FOXM1. Elevated FOXM1 levels were recently reported in aggressive meningiomas (19) (as well as other cancers (34)), but our findings suggest that this upregulation is secondary to the loss of DREAM complex-mediated repression. Our findings suggest that FOXM1 and MYBL2 act as co-activators of the DREAM complex rather than as independent transcription factors.

Of note, 79% of type C tumors showed loss of both 1p and 22q. All the WHO grade III tumors in our sample (which were all recurrences) show this double loss, but two of the patients had multiple resections of WHO grade II tumors. Our molecular classification, by contrast, placed all these patients’ tumors, from original to last, in type C. Moreover, that our type C tumors, the most aggressive type, included all WHO grades underscores the importance of developing robust molecular profiles to supplement histopathology.

Our data suggest that testing for loss of even just these two chromosomes could provide a valuable biomarker for the risk of recurrence despite complete resection. In addition, it is worth noting that copy number variation alone may prove to be sufficient in distinguishing these three tumor types, and certainly would be a simple test to carry out clinically. To solidify this correlation and its diagnostic value, future studies should evaluate these genomic loci in much larger samples.

The pressing need in the meningioma field is to understand the biology that differentiates aggressive meningiomas from less aggressive ones so that we may start dissecting the pathways that drive pathogenesis and establish the first step towards developing adjuvant therapies. Our three molecular types differ clinically and biologically and correlate with the clinical course better than the WHO classification. We continue to follow up on our patient population with the expectation that more data will yield further insight. Along similar lines, much larger cohorts will be needed to refine the molecular profiles (both genomic data and RNA expression) to a clinically translatable signature, to better understand meningioma biology and improve prognostication of the most difficult meningiomas.

## Methods

### Sample Selection and Preparation

We obtained 161 primary tumor tissue (fresh frozen) samples from 141 patients who were treated at Baylor College of Medicine (BCM). All patients provided written informed consent, and tumor tissues were collected under an IRB approved protocol at BCM by the Human Tissue Acquisition and Pathology Core. All meningiomas were initially signed out by one of two neuropathologists (K.H. or J.C.G.) and were graded based on the 2016 WHO guidelines. MIB-1 index was calculated by determining the percentage of meningioma cell nuclei positive for Ki-67 staining. We used blood DNA as a reference for detecting somatic tumor mutations. We performed RNA sequencing on 161 tumors. One tumor sample was noted to have a NAB2-STAT6 gene fusion, that, based on the 2016 WHO guidelines (47), is now diagnostic for hemangiopericytoma/solitary fibrous tumor. Upon independent review by our neuropathologist, the patient was excluded from our analysis. We thus analyzed 160 meningiomas from 140 patients: 121 benign (WHO grade I), 32 atypical (WHO grade II) and 7 malignant (WHO grade III) meningiomas. One-hundred and twenty-eight of these samples had adequate DNA for whole-exome sequencing. Only representative fresh-frozen blocks with estimated purity of ≥95% were selected for DNA and RNA extraction from 20-30 mg of tumor tissue using TRIzol (Thermo Fisher Scientific, MA) according to the manufacturer’s protocol. Normal DNA was extracted from 1 ml of whole blood stored in PAXgene blood DNA tubes using the PAXgene Blood DNA Extraction Kit (Qiagen, CA) according to the manufacturer’s protocol.

### Patient Data and Characteristics

Under the aegis of a BCM IRB-approved protocol, we reviewed the following data: patient age at surgery, gender, race, tumor size, tumor location, pre-operative embolization, extent of resection, histologic grade by WHO guidelines, MIB-1 index, and presence of brain invasion. Diagnostic imaging was re-reviewed to define tumor location, extent of resection and presence/date of local recurrence. Local recurrence after gross total resection was defined as local development of any contrast enhancement on subsequent brain imaging. Local recurrence after subtotal resection was defined as measurable growth of residual tumor. Vital status of the patient was obtained from search of the electronic medical record. A summary of clinical information is available in Table 1.

**Table 1:** Clinical features of patients and their primary meningiomas. Note that this table does not include recurrences (see Table S1).

|  | Discovery set |  |  |  |  | Validation set |  |  |  |
| --- | --- | --- | --- | --- | --- | --- | --- | --- | --- |
| Characteristic | All tumors<br>(no.=97) | Type A<br>(no.=35) | Type B<br>(no.=30) | Type C<br>(no.=32) | p-value<br>btwn types | All tumors<br>(no.=48) | Type A<br>(no.=23) | Type B<br>(no.=10) | Type C<br>(no.=15) |
| sex, no. of patients (%) |  |  |  |  |  |  |  |  |  |
| male | 36 (38) | 12 (34) | 7 (23) | 18 (56) | **0.0240** | 14 (30) | 3 (13) | 6 (60) | 6 (40) |
| Female | 60 (62) | 23 (66) | 23 (77) | 14 (44) |  | 32 (70) | 20 (87) | 4 (40) | 9 (60) |
| median age (years) at surgery (range) | 60<br>(27-81) | 62<br>(33-79) | 58.5<br>(27-81) | 58.5<br>(33-78) | 0.465 | 59<br>(18-81) | 58<br>(21-78) | 63.5<br>(26-81) | 63<br>(18-77) |
| Location (%) |  |  |  |  |  |  |  |  |  |
| Anterior Cranial Fossa | 15 (16) | 13 (37) | 2 (7) | 0 (0) | **0.0001** | 7 (15) | 7 (30) | 0 (0) | 0 (0) |
| Middle Cranial Fossa | 2 (2) | 2 (6) | 0 (0) | 0 (0) | 0.1930 | 2 (4) | 2 (9) | 0 (0) | 0 (0) |
| Sphenoid Wing | 16 (16) | 9 (26) | 5 (17) | 2 (6) | 0.1930 | 9 (19) | 5 (22) | 2 (20) | 2 (13) |
| Parafalcine | 15 (15) | 1 (3) | 7 (23) | 7 (22) | 0.1800 | 6 (13) | 1 (4) | 1 (10) | 4 (27) |
| Petroclival | 7 (7) | 1 (3) | 5 (17) | 1 (3) | 0.1930 | 2 (4) | 2 (9) | 0 (0) | 0 (0) |
| Clival | 4 (4) | 1 (3) | 2 (7) | 1 (3) | 0.9030 | 9 (19) | 3 (13) | 4 (40) | 2 (13) |
| Frontal | 14 (14) | 5 (14) | 1 (3) | 8 (25) | 0.1800 | 7 (15) | 1 (4) | 2 (20) | 4 (27) |
| Occipital | 5 (5) | 0 (0) | 0 (0) | 5 (16) | **0.0330** | 2 (4) | 2 (9) | 0 (0) | 0 (0) |
| Parietal | 6 (5) | 1 (3) | 2 (7) | 3 (9) | 0.7290 | 1 (2) | 0 (0) | 0 (0) | 1 (7) |
| Temporal | 6 (6) | 1 (3) | 2 (7) | 3 (9) | 0.7290 | 0 (0) | 0 (0) | 0 (0) | 0 (0) |
| Tentorial | 1 (1) | 1 (3) | 0 (0) | 0 (0) | 0.4850 | 0 (0) | 0 (0) | 0 (0) | 0 (0) |
| Intraventricular | 2 (2) | 0 (0) | 0 (0) | 2 (6) | 0.1930 | 1 (2) | 0 (0) | 0 (0) | 1 (7) |
| Cerebellum | 2 (2) | 0 (0) | 2 (7) | 0 (0) | 0.1930 | 0 (0) | 0 (0) | 0 (0) | 0 (0) |
| Spine | 2 (2) | 0 (0) | 2 (7) | 0 (0) | 0.1930 | 1 (2) | 0 (0) | 1 (10) | 0 (0) |
| WHO Grade (%) |  |  |  |  |  |  |  |  |  |
| Grade I | 77 (79) | 35 (100) | 24 (80) | 18 (56) | **0.0020** | 39 (81) | 23 (100) | 8 (80) | 8 (53) |
| Grade II | 20 (21) | 0 (0) | 6 (20) | 14 (44) |  | 9 (19) | 0 (0) | 2 (20) | 7 (47) |
| Median MIB1 Index (range) | 3.1<br>(0.5-40) | 2.5<br>(0.5-18.5) | 3.5<br>(0.5-31.5) | 6.3<br>(1-40) | **0.0040** | 2.6<br>(1-32.7) | 2.2<br>(1-5.5) | 5<br>(2.5-16.5) | 8.2<br>(1-32.7) |
| Extent of Resection (%) |  |  |  |  |  |  |  |  |  |
| Gross Total Resection | 76 (79) | 25 (71) | 27 (90) | 24 (75) | 0.4820 | 38 (79) | 16 (70) | 9 (90) | 13 (87) |
| Subtotal Resection | 20 (21) | 10 (29) | 3 (10) | 7 (22) |  | 10 (21) | 7 (30) | 1 (10) | 2 (13) |
| Unknown | 1 (1) | 0 (0) | 0 (0) | 1 (3) |  | 0 (0) | 0 (0) | 0 (0) | 0 (0) |
| Median Follow-up (months; range) | 28<br>(0-91) | 26<br>(8-86) | 25<br>(1-83) | 31<br>(0-91) | 0.5720 | 11.5<br>(0-20) | 5<br>(0-17) | 3<br>(0-17) | 4<br>(0-20) |
| Death (%) | 1 (1) | 0 (0) | 0 (0) | 1 (3) | 0.3580 | 1 (2) | 0 (0) | 1 (11) | 0 (0) |

The breakdown of patients and profiled recurrences is as follows: 126 patients had only one tumor (126 tumors); five patients had two distinct tumors (for a total of 10 tumors); six patients had one recurrent tumor (12 tumors); one patient had a recurrence with 2 separate tumor masses (3 tumors); one patient had 4 sequential recurrences (5 tumors); one patient had two sequential recurrences, where the second recurrence produced two distinct masses (4 tumors). This yielded a total of 160 tumors (126 + 10 + 12 + 3 + 5 + 4).

To look at the data another way, the discovery cohort contained 97 tumors from 95 patients on whom we operated between 2011 and 2017, including two patients that had two primary tumors. The validation set contained 48 tumors from 47 patients on whom we operated between 2017 and 2018, including one patient with two primary tumors. Two patients had a primary tumor in the discovery set and had a second primary tumor that ended up in the validation set.

### Antibodies

Western blot (overnight incubation with a 1:5,000 dilution): anti-EED (chicken, GTX14294, GeneTex, TX), anti-SUZ12 (D39F6, rabbit, 3737S, Cell Signaling Technology, MA), anti-Vinculin (hVIN, mouse, V9131, MilliporeSigma, MA), anti-FOXM1 (rabbit, GTX102126, GeneTex, TX), anti-MYBL2 (rabbit, GTX77893, GeneTex, TX), anti-RBL2 (D9T7M, rabbit, 13610, Cell Signaling Technology, MA), anti-Mouse-HRP (1:50:000, 715-035- 150, Jackson ImmunoResearch Labs, PA, RRID:AB_2340770), anti-Rabbit-HRP (1:20,000, 170–5046 Bio-Rad/AbD Serotec, CA, RRID:AB_11125757), anti-Chicken-HRP (1:2,000, NBP1-74785, Novus Biologicals, CO). Co-IP: anti-EZH1 (rabbit, 2 μg per IP, GTX108013, GeneTex, TX), anti-LIN37 (rabbit, 5 μg per IP, GTX44925, GeneTex, TX).

### Co-Immunoprecipitation

PRC2 immunoprecipitations were carried out with 10 mg of tissue in 200 μl of PRC2 lysis buffer [50 mM HEPES (pH 7.0), 250 mM NaCl, 0.1% Nonidet P-40, 5 mM EDTA, freshly added: 0.5 mM dithiothreitol, 1 mM PMSF, 1X Xpert Phosphatase Inhibitor and 1X Xpert Protease Inhibitor Cocktail (P3200 and P3100, GenDEPOT, TX)]. DREAM immunoprecipitations were carried out with 50 mg of tissue in 200 μl of DREAM lysis buffer [20 mM Tris (pH 7.5), 420 mM NaCl, 1.5 mM MgCl2, 1 mM EDTA, 5% glycerol, freshly added: 1 mM dithiothreitol, 1 mM PMSF, 1X Xpert Phosphatase Inhibitor and 1X Xpert Protease Inhibitor Cocktail]. After tissue disruption via sonication (three rounds with 3, 4 and 5 pulses respectively at 20% duty cycle), lysates were cleared via centrifugation for 20 minutes at 21,000 RCF at 4°C and transferred to siliconized tubes. The antibody was added and after a 2- hour incubation at 4°C on a rotor, 40 μl of agarose beads were added for another 30 minutes. Antibody-bead complexes were washed five times in their respective buffers and subject to standard western blot analysis using 1% input and 50% eluates.

### Cell culture and qRT-PCR

Arachnoid cells were immortalized using a lentivirus harboring the SV40 large T antigen [pBABE-puro SV40 LT was a gift from Thomas Roberts (Addgene, RRID:Addgene_13970) as previously described (48). NF2 haplotype was validated using qRT-PCR using a dilution series of DNA from 293T (NF2 wildtype) and arachnoid cells. 293T cell line was purchased from ATCC (CRL-3216, Manassas, VA). Cells were found to be negative for mycoplasma contamination. Cell lines were cultured as adherent cells in DMEM containing 10% FBS and antibiotics using standard cell culture practices (Geraghty et al., 2014). 293T or arachnoid cells were transfected using Lipofectamine 3000 (Thermo Fioscher, Carlsbad, CA)using the following constructs: hEZH1-GFPpcDNA3 (aa1-747), EZH1_deltaSET-GFPpcDNA3 (aa1-512), hLIN37-GFPpcDNA3 (aa1-243), hLIN37_CD1/2-GFPpcDNA3 (aa1-243) (41), hLIN52-GFPpcDNA3 (1-116) and Lin52_S28A-GFPpcDNA3) (42). After 48 hours of culture, total RNA was isolated using TRIzol, subject to reverse transcription and qRT-PCR. Primer information is available upon request.

#### Data deposition

Raw and processed data will be available in the GEO repository upon publication.

See extended methodological details in SI Appendix

## Acknowledgements

We would like to thank the patients, without whom this study would not be possible. We would like to thank Chad Shaw, PhD for reviewing the paper to ensure proper bioinformatics/statistical methodology was used. Portions of this study were funded by the Roderick D. MacDonald Fund, the Jan and Dan Duncan Neurologic Research Institute at Texas Children’s Hospital, and the Hamill Foundation. AJP is supported by a K08 award from the NINDS (K08NS102474). HYZ is supported by the Howard Hughes Medical Institute. The Human Tissue Acquisition & Pathology (HTAP) Core at Baylor College of Medicine is funded through the P30 Cancer Center Support Grant (NCI-CA125123). We thank Vicky Brandt for insightful comments on the manuscript; Kathy Relyea for the illustration in Fig. 2C; and Dima Suki for providing advice on statistical analysis.

## Contributions

A.J.P. conceived the main study idea; A.J.P, T.J.K. and H.Y.Z. designed and/or performed experiments, analyzed data, and wrote and revised the manuscript. Y.W.W., R.A.O. and Z.L. performed and interpreted computational analyses. J.F.M. performed statistical analysis. J.P.R., M.O. and S.S. performed sample preparation of patient tissue. M.C., L.X., D.M.M., H.D., and D.A.W. performed next generation sequencing experiments. K.A.H. and J.C.G. performed histopathological classification. A.J.P., S.P.G., and D.Y. provided patient samples. A.J., D.A.W., G.R., D.Y., S.E.P., and H.Y.Z. provided feedback on study design, reviewed the data, and revised the manuscript. All authors gave final approval.

