## Supplement information for "Molecular profiling predicts meningioma recurrence and reveals loss of DREAM complex repression in aggressive tumors"

**This PDF file includes**

Figures S1 to S2

Legends for Datasets S1 to S4

Extended Methodological Details

SI References

**SI Figures:**

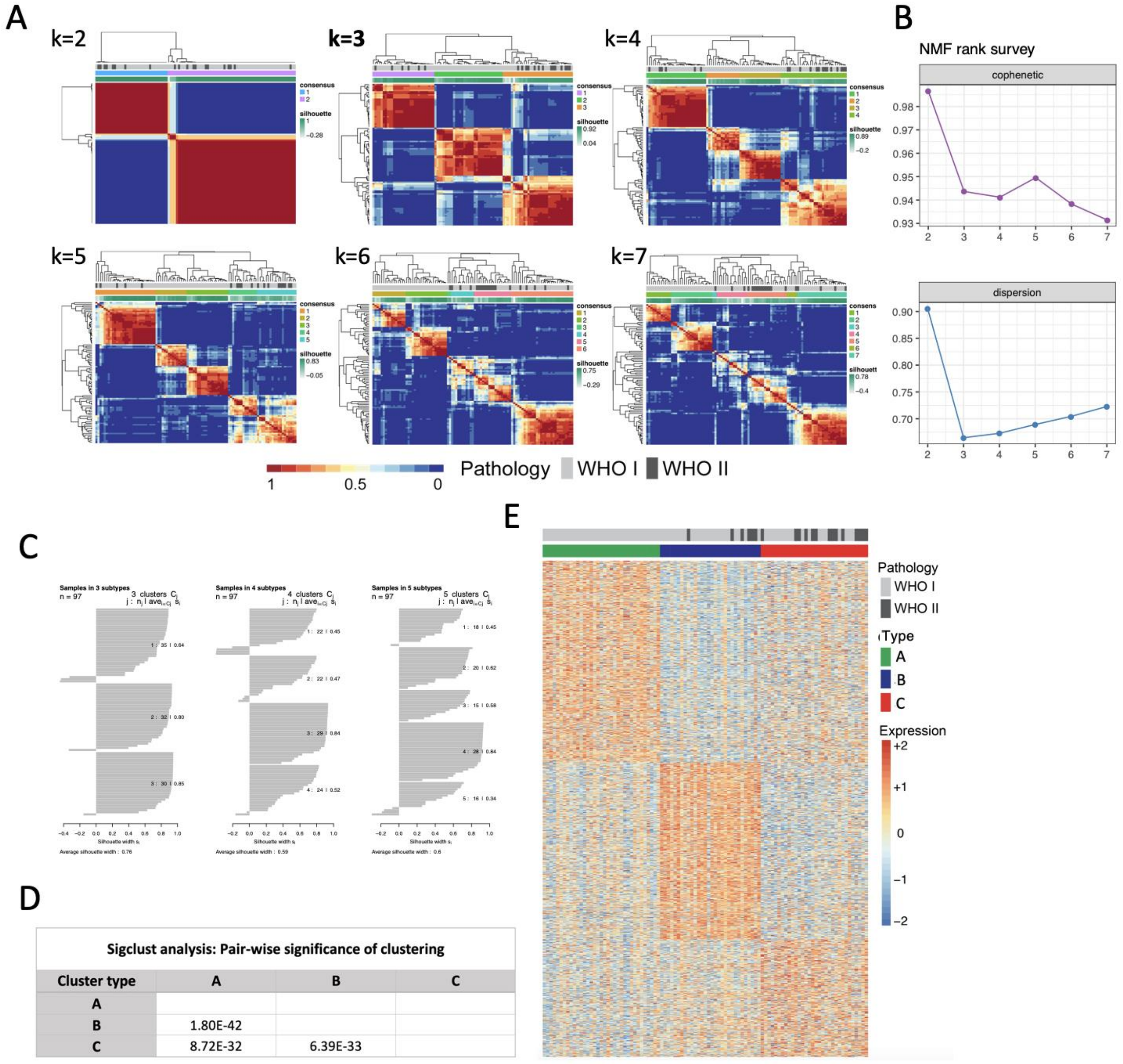


| Cluster type | A | B | C |
|---|---|---|---|
| A | | | |
| B | 1.80E-42 | | |
| C | 8.72E-32 | 6.39E-33 | |

**Fig. S1 NMF for rank k=2 to k=7 on the 1500 most variable genes.** (**A**) Heatmaps on the consensus matrix shows the average connectivity of the 97 samples from the 1000 runs for each rank. (**B**) Cophenetic correlation coefficients and dispersion associated with clusters observed from each rank. (**C**) Silhouette analysis on clusters observed from rank 3 to 5 shows that k=3 is the optimal number of clusters as all clusters give average silhouette width of greater than 0.6, while there exist clusters lower than 0.5 for other ranks. (**D**) Significance table of clusters pair-wise comparisons. (**E**) Expression heatmap of the top 1,500 most variable genes in the discovery set. Type Ais labelled in green; type B, blue; type C, red.

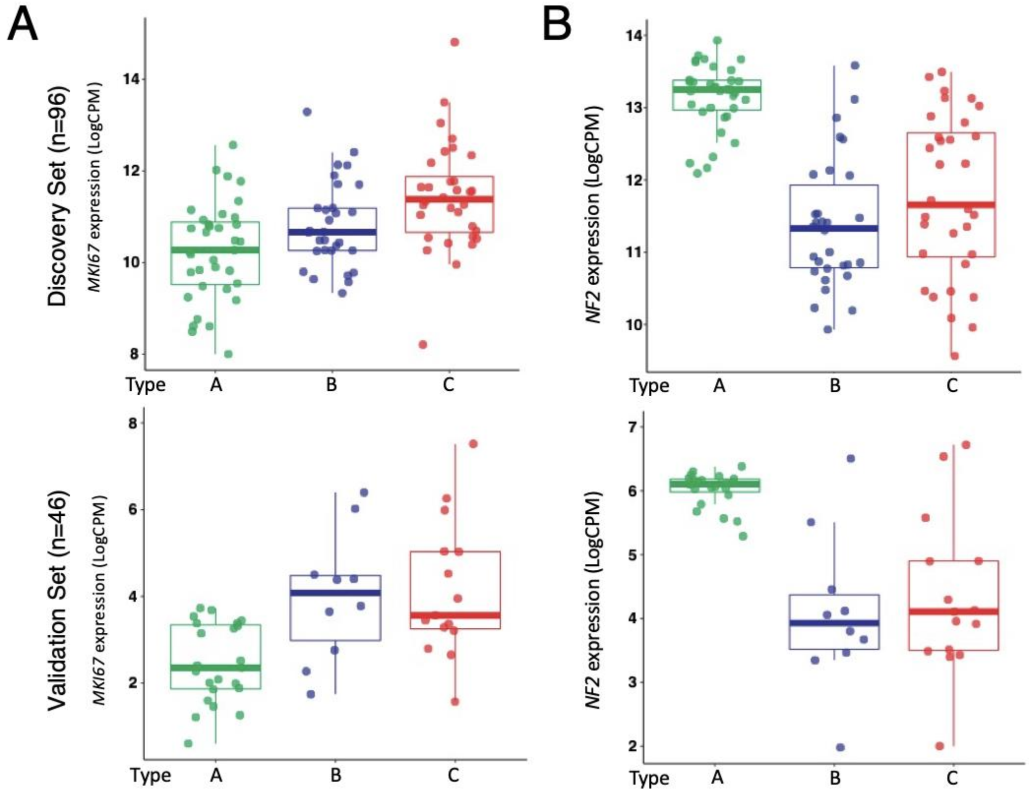


**Fig. S2 MKI67 and NF2 expression level by type** (**A**) Boxplot demonstrating MKI67 expression for types A to C in the discovery set (upper panel) and validation set (lower panel). (**B**) Boxplot showing NF2 expression in each of the three types in the discovery set (upper panel) and validation set (lower panel). NR2 expression between types B and C is not statistically different.

**SI Datasets:**

| | |
|---|---|
| Table S1 | Detailed tumor data |
| Table S2 | NF2 mutations in discovery and validation sets |
| Table S3 | Differentially expressed gene lists |
| Table S4 | GSEA analysis of expression types |

### Extended Methodological Details

*Data reporting*

No statistical methods were used to predetermine sample size. The experiments were not randomized, and the investigators were not blinded to allocation during experiments and outcome assessment.

*Next generation sequencing and analysis*

All protocols and analyses were performed by the Human Genome Sequencing Center of Baylor College of Medicine (BCM-HGSC, https://www.hgsc.bcm.edu) according to their standard operation procedures.

*DNA sequencing*

DNA libraries were constructed according to the manufacturer's protocol with modifications. Briefly, 0.5 μg DNA was sheared into fragments of 200–300 base pairs in a Covaris plate with E210 system (Covaris, MA) followed by end-repair, A-tailing and ligation of Illumina multiplexing PE adaptors. Pre-capture Ligation Mediated-PCR (LM-PCR) was performed using the Library Amplification Readymix containing KAPA HiFi DNA Polymerase (Kapa Biosystems, MA). Universal primer IMUX-P1.0 and IMUX-P3.0 were used in PCR amplification. Purification was performed with Agencourt AMPure XP beads (Beckman Coulter, CA) after enzymatic reactions. Following the final XP beads purification, quantification and size distribution of the pre-capture LMPCR product was determined using the LabChip GX electrophoresis system (PerkinElmer, MA) and gel analysis using AlphaView SA v3.4 software. For exome capture, four pre-capture libraries were pooled together. The pooled libraries were then hybridized in solution to the HGSC VCRome 2.1 design (42Mb, Roche NimbleGen, WI). Post-capture LM-PCR amplification was performed using the Phusion High-Fidelity PCR Master Mix. After the final AMPure XP bead purification, quantity and size of the capture library was analyzed using the Caliper LabChip GX electrophoresis system. The efficiency of the capture was evaluated by performing a qPCR-based quality check on the enrichment level of four standard NimbleGen control loci. All sequencing runs were performed in PE mode on Illumina HiSeq2000 platform.

*Bioinformatics analyses for whole exome-sequencing data*

Initial sequence analysis was performed using the HGSC Mercury analysis pipeline, from which the sequence reads of each sample were mapped to GRCh37 Human reference genome using the Burrows-Wheeler aligner (BWA, RRID:SCR_010910) and then recalibrated and realigned by GATK (RRID:SCR_001876). Mutations called by BCM-HGSC were generated by the standard cancer analysis pipeline, CARNAC (Consensus and Repeatable Nucleotide Alterations in Cancer), as described previously (1, 2).

*RNA sequencing*

Genomic RNA was quantified by Picogreen (Thermo Fisher Scientific, MA) and its quality was assessed using a 2100 Bioanalyzer (Agilent, CA). RNA was selected for sequencing only if the RIN value was >6. RNA samples (1 μg) were fragmented, converted into double-stranded cDNA and then proceeded to library prep using TrueSeq RNA Sample Preparation (Illumina, San Diego, CA) according to the manufacturer protocol using a 15-cycle amplification. After PCR primers removal using Agencourt AMPure PCR Purification kit (Beckman Coulter, CA), the libraries were analyzed and quantified on TapeStation (TapeStation Instrument, RRID:SCR_014994) using the DNA High Sensitivity kit (Agilent, CA) to verify correct fragment size and to ensure the absence of extra bands. Equimolar amounts of DNA were pooled for capture (8 samples per pool) and verified by TapeStation. The captured libraries were sequenced on a HiSeq 4000 (Illumina HiSeq 4000 System, RRID:SCR_016386) on a version 3 TruSeq paired end flowcell according to manufacturer's instructions at a cluster density between 700 and 1000 K clusters/mm$^2$. The resulting BCL files containing the sequence data were converted into ".fastq.gz" files and individual libraries within the samples were demultiplexed using CASAVA 1.8.2 (CASAVA, RRID:SCR_001802) with no mismatches. All regions were covered by >20 reads.

*Bioinformatics analyses for RNA sequencing data*

Raw reads from the RNA-seq samples were processed using an in-house pipeline that uses TopHat2 (RRID:SCR_013035) for read alignment; FastQC (RRID:SCR_014583) and RSeQC (RRID:SCR_005275) for read and alignment quality assessment; HTSeq (RRID:SCR_005514) for expression count; and GATK (RRID:SCR_001876) for variant calling. The reads were aligned to GRCh37 human reference genome and mapped to the human transcriptome according to UCSC gene annotations. The RNA-seq read counts for genes were normalized and transformed into log-counts using the voom method (variance modeling at the observational level) (3) implemented in R package limma (4). The normalized counts were then batched corrected using ComBat method implemented in R package sva (5). Batch-corrected expression data were used for subsequent subtype analysis using R software package NMF (https://cran.r-project.org/web/packages/NMF/index.html). Expression heatmaps were generated using Pheatmap (RRID:SCR_016418). Differential gene expression analysis of genes between the tumor types identified was carried out using the limma package on batch-corrected expression data.

To compare the discovery set with the validation sets, a list of 3,484 genes that were expressed in all three data sets was identified. Using this gene set, we were able to compare the three subtypes in a pairwise manner and identified differentially expressed genes (FDR of 1% and fold change >= 1.5) using the limma package. To verify the gene signature in the validation datasets, we first transformed the gene counts in the training and validation datasets to z-scores. We then built a Random Forest classifier using the gene signature and used it to predict the subtype classes for the samples in the validation datasets. Heatmaps for the validation datasets were produced using the assigned labels from the RF and similar gene ordering as in the training dataset heatmap. Heatmaps were implemented using the R package pheatmap.

*Statistical analysis*

For all analyses, a p-value<0.05 was considered significant. Recurrence-free survival (RFS) analysis and subsequent survival data visualization were carried out in R, using survival and survminer (https://CRAN.R-project.org/package=survminer) packages, respectively. ANOVA and Chi-square tests were used to compare clinical variables between types. We used R to fit a generalized linear model (GLM) with a poisson link function (R function glm) to compare the rates of tumors by type and location. Main effects and interactions were tested using an analysis of deviance (chi-square), provided by the car package (https://CRAN.R-project.org/package=car). After a significant type-by-location interaction, we used further chisquares tests to determine tumor type differences within each location. False discovery rate correction was used to account for Type I error inflation due to multiple comparisons (16 locations). To compare type-by-location patterns between our discovery and validation datasets, we used the Bayesian Information Criterion (BIC) to compare two GLMs: one GLM contained interactions with dataset, the other model contained only a main effect of dataset (to account for differences in sample size). The model with the lower BIC provides the better fit to the data, with a BIC difference of 10 used as a threshold for decisive evidence in favor of a given model.

Chi-square tests were performed to compare proportions of gender and frequency of chr22q loss, chr1p loss, and TERT promoter mutations across the three types. To compare the frequency of NF2 mutations, a Chi-square test between type B and C tumors was performed since there were no NF2 mutations in type A tumors.

*Gene Set Enrichment Analyses*

One over-expressed gene list per tumor type was generated using RNA-seq expression data that had a logFoldChange >1 with a false discovery rate <0.001 (Table S3). Gene Set Enrichment Analyses (GSEA, RRID:SCR_003199) were performed as described elsewhere (31) using curated gene sets (C2). All gene sets with an overlap of 15% or more percent were considered in the analysis and can be found in Table S4. The following gene sets from the Broad Institute (https://software.broadinstitute.org/gsea/) were used: PRC2 target genes (combination of M9898, M7617, M8448), DREAM target genes (M149), Cell cycle genes (M7963). The HOX genes that are known to be repressed by the PRC2 complex were generated by identifying the intersection the homeodomain proteins (47) with the following GSEA gene lists: M7617, M10371, M1938, M8448, M9898. This identified 163 HOX genes.
